# Distinct pathogenic influence of anti-HMGCR+ and anti-SRP+ immune-mediated necrotizing myopathy autoantibodies on engineered muscle function

**DOI:** 10.1101/2024.08.26.609771

**Authors:** Heta Lad, Ernest Myguel Esteban, Yekaterina Tiper, Alexandrine Mahoudeau, Zhuoye Xie, Berenice Tendrel, Yves Allenbach, Olivier Benveniste, Penney M. Gilbert

**Author notes:** Address correspondence to Penney M. Gilbert, PhD: 160 College Street, M5S 3E1, Toronto, Canada. Address correspondence to Olivier Benveniste, MD, PhD: MITIC U974, 3ème étage, 105 Boulevard de l’Hôpital CHU Pitié-Salpetrière, 75651, Paris, France. These authors contributed equally (Heta Lad, Ernest Myguel Esteban). These authors contributed equally (Alexandrine Mahoudeau, Zhuoye Xie).

## Abstract

Immune-mediated necrotizing myopathy (IMNM) is a subgroup of idiopathic inflammatory myopathies associated with anti-signal recognition particle (SRP) or anti-3-hydroxy-3-methylglutaryl-CoA reductase (HMGCR) autoantibodies. However, the demonstration of a direct pathogenic effect of IMNM patient autoantibodies on skeletal muscle contractile force, independent of the downstream activation of the complement pathway, has yet to be reported. Thus, the goal of this study was to leverage a custom 3D-human skeletal muscle microtissue (hMMT) culture platform, that enables muscle cell contractile apparatus maturation and the analysis of contractile function, to evaluate the direct effect of total immunoglobulins (IgGs) isolated from IMNM patients with amplification of anti-SRP^+^ or anti-HMGCR^+^ autoantibodies. hMMTs capable of force generation were treated with total IgGs, isolated from 3 SRP+ and 3 HMGCR+ patients plasma, and delivered in complement inactivated media for 4 days. hMMT health was then evaluated by quantifying the peak force and contraction kinetics in response to electrical field stimulation and by performing histological analysis of sarcomere and myotube structures. Treating hMMTs with total IgGs from anti-HMGCR^+^ patients resulted in a decline in tetanus contractile force, though sarcomere Z-line architecture analysis revealed no significant influences on sarcomere organization. hMMT treatment with total IgGs from anti-SRP^+^ patients induced muscle atrophy, observed via significantly smaller myotube diameter, but this did not translate to a decline in contractile function. This study demonstrates that anti-SRP and anti-HMGCR autoantibodies exert direct, but distinct influences on IMNM-associated skeletal muscle pathogenesis, which may inform IMNM therapy development.

## 1.1 Introduction

Idiopathic inflammatory myopathies (IIM), also known as myositis, are a heterogenous family of autoimmune disorders where muscle weakness and myalgia are classical clinical manifestations developing gradually over a period of weeks to months, or even years, and frequently resulting in severe impairment [1–5]. Other organ systems may be additionally affected, and prompt life-threatening complications [1–5]. Advances in IIM studies have led to a classification of subtypes based on differences in histopathology or clinical manifestations, one of which is immune-mediated necrotizing myopathy (IMNM) [2,3].

IMNM is the most severe and disabling form of myositis and accounts for 35% of all IIM cases [1,2,4]. This subgroup of IIMs is characterized by rapidly progressive muscle weakness and substantially elevated creatine kinase (CK) levels, often requiring early, aggressive immunotherapy [1,4,5]. To date, the presence of two autoantibodies (aAb) has been associated with IMNM; anti-signal recognition particle (SRP) or anti-3-hydroxy-3-methylglutaryl-CoA reductase (HMGCR) aAbs. Anti-SRP^+^ IMNM can be severe and progress rapidly, or it can progress slowly, mimicking limb-girdle muscular dystrophy [6,7]. Anti-HMGCR^+^ IMNM is also severe and may occur during or after statin exposure [8]. Both anti-SRP and anti-HMGCR aAb titers have been shown to correlate highly with CK levels and muscle strength [9,10]. Evidence of aAb involvement in IMNM has been shown in studies whereby anti-SRP and anti-HMGCR aAbs recognized and bound to the target autoantigens exposed in the sarcolemma of myofibers leading to activation of the classical complement pathways, which resulted in muscle damage [11]. Interestingly, a phase II clinical trial testing a complement inhibitor, zilucoplan, did not improve the outcome measures in patients with IMNM, suggesting the possibility that complement activation might be secondary to skeletal muscle damage [12]. Furthermore, the expression of SRP and HMGCR is ubiquitous, rather than muscle specific. Thus, pathophysiological mechanisms leading to severe muscle impairments in IMNM remain to be elucidated.

CAR-T Cell therapy for autoimmune disorders is emerging as a breakthrough in the field[13]. Indeed, CAR-T Cells targeting CD19 B cells [13] or B cell maturation antigen [14] have shown impressive results in treating idiopathic myositis patients, and anti-SRP IMNM patients more specifically, in the latter case. The success of targeting antibody-producing B cells for destruction emphasizes the critical need to understand the pathogenic role of Abs in IMNM to guide next generation therapeutic cell products. A direct pathogenic effect of aAbs may initially seem unlikely since SRP and HMGCR are intracellular targets. However, genetic HMGCR deficiency leads to muscle dystrophy [15], and although only 4% of muscle fibers show necrosis in anti-HMGCR myositis muscle biopsies [11], patients display severe weakness. Further, myositis-specific antibodies have been shown to penetrate muscle fibers *in vitro* [16], opening the plausibility of direct impacts of anti-SRP and anti-HMGCR auto-antibodies on muscle fiber function.

While there is limited modeling of IMNM, some work has been conducted to investigate the pathogenicity of SRP and HMGCR aAbs in animal models and with traditional 2D skeletal muscle culture systems. In a study where mice were injected with plasma or purified IgGs from anti-SRP^+^ and anti-HMGCR^+^ patients, a decline in mouse *gastrocnemius* muscle and grip strength was observed[17]. However, in the context of immunocompetent mice, the occurrence of a xenogenic response to injected human IgGs prevents the study of long-term disease in this model. Also, isolating direct influences of pathogenic aAbs on skeletal muscle in an *in vivo* setting is difficult since they cannot be easily decoupled from immune influences. As an alternative, 2D skeletal muscle cultures reduce complexity and enable investigation of the direct effects of aAbs on muscle health, independent of the activation of downstream complement pathways. It has been shown that anti-SRP and anti-HMGCR aAbs impair myoblast fusion and induce myotube atrophy *in vitro* [18]. However, a 2D platform is not well suited to evaluations of skeletal muscle function since contractile myotubes often delaminate from the culture dish and the absence of a three-dimensional extracellular matrix environment stunts contractile apparatus maturation [19]. Thus, while capable of capturing a direct myotube atrophying effect, 2D culture systems are fundamentally limited in demonstrating how muscle function is directly affected by anti-SRP and anti-HMGCR aAbs.

To address these gaps, we implemented a three-dimensional (3D) skeletal muscle cell culture system that allows the modeling of more physiologically and pathologically relevant processes *ex vivo* [20,21]. Specifically, the (Myo) microTissue Array device To Investigate forCe (MyoTACTIC) culture device enables the bulk production of 3D human skeletal muscle microtissues (hMMTs) within a 96-well plate format wherein the deflection of flexible vertical posts enables *in situ* quantification of tissue strength and calcium transient behaviors [20–22]. Leveraging this system, we treated hMMTs with total IgGs purified from the plasma of anti-SRP^+^ and anti-HMGCR^+^ patients to investigate whether these aAbs induce a direct pathogenic effect on muscle cell morphology and force-generating capacity of hMMTs.

## 1.2 Materials and Methods

### 1.2.1 Patient samples and total IgG extraction

IMNM patient plasma was collected during plasmapheresis and stored in our biobank MASC (Myositis, DNA, Serum, Cells, ClinicalTrials.gov Identifier: NCT04637672). Approvals by the Local Ethics Committee (CPP Ile De France VI (2013-12-19), CCTIRS (N°14.323) and CNIL (915139)) were obtained to conduct the studies described herein. Written informed consent was obtained from each patient. Pharmaceutical grade OCTAPLAS LG® human plasma (Group A, Lot M726A9521) was purchased to extract healthy control IgG (OctoPharma AB). Before IgG extraction, samples were diluted (1:1) in BupH^TM^ phosphate buffered saline (Cat. #28372, Thermo Scientific) and then centrifuged at 3200 g for 20 minutes to equilibrate pH and avoid column blockade. To extract total IgG from each sample, protein G spin columns were used by following the manufacturer instructions (Cat. #89957, Thermo Scientific). After extraction, total IgG eluate was concentrated using 30 kDa Amicon filters (Cat. #UFC9030; Millipore) and dialyzed once with Dulbecco’s Phosphate Buffered Saline (D-PBS; Cat. #311-415-CL; Wisent Bioproducts), and then concentrated again to obtain a stock concentration between 45 and 55 mg/ml. Total IgG concentration was quantified using NanoDrop 1000 Spectrophotometer before storage at -20°C. Antigen recognition by patient antibodies was tested using a kit (for anti-SRP; Cat. #PMS12DIV-24; D-tek), or an automated machine (for anti-HMGCR; Cat. #701333, Inova Diagnostics) by following the supplier instructions in Dr. Charuel’s immunology laboratory (Pitié Salpêtrière Univeristy Hospital, Paris France). The total IgG concentration extracted for each sample and its target recognition is provided in **Supplementary Table 1**.

### 1.2.2 MyoTACTIC culture platform fabrication

The polydimethylsiloxane (PDMS; Sylgard^TM^ 184 silicone elastomer kit; Dow Corning) MyoTACTIC 96-well culture platform was used to generate human skeletal muscle micro-tissues (hMMTs). In this platform, each well contains an oval pool with two flexible PDMS micro-posts on either side of the pool. These posts provide uniaxial tension to remodeling hMMTs and enable the non-invasive quantification of hMMT contractile function. The platform was fabricated exactly as we previously described [23,24]. All features of the plate were cast in a single step from a reusable, negative polyurethane (PU; Smooth-Cast® 310 liquid plastic; Smooth-On) mold. PDMS culture plates were prepared for muscle cell culture by first sonicating MyoTACTIC portions for 20 minutes in isopropanol (Cat. 8600), after which they were rinsed in ddH_2_O and set to evaporate in a curing oven for 15 minutes. MyoTACTIC portions were autoclaved for final sterilization before use.

### 1.2.3 Human immortalized myoblast cell line maintenance

The AB1167 (Healthy) and AB1071 (DMD) human-immortalized myoblast cell lines used in the fabrication of hMMTs for this study were established at the Myoline platform of the Institut de Myologie (Paris, France) [25]. The myoblast line was expanded, as previously described [24,26], to produce the requisite number of cells for hMMT seeding. A summary of all media and solutions used for cell line expansion can be found in **Supplementary Table 2**.

### 1.2.4 hMMT seeding and total IgG treatment

3D hMMTs were generated in the MyoTACTIC platform from AB1167 (Healthy) and AB1071 (DMD) immortalized cells exactly as we previously described [24,26]. Following seeding, constructs were incubated in hMMT after seeding growth media (Day -2). Two days later, the culture media was exchanged for differentiation media (DM; Day 0) with half media changes carried out every other day until day 8 of differentiation. On day 8, the standard DM containing horse serum (HS; Cat. #16050114; Gibco) was removed and replaced with DM containing either HS, heat-inactivated (HI) HS, HI HS with cerivastatin (25 nM or 75 nM in ddH_2_O; Cat. #SML0005; Sigma) or HI HS with 1 mg/ml of healthy, anti-SRP or anti-HMGCR total IgGs. Half of the media was refreshed with the respective treatment on day 10 and end-point analyses were performed on day 12 of differentiation. Total IgG from three patients identified to be producing anti-SRP autoantibodies and three patients producing anti-HMGCR autoantibodies served as biological replicates. Three technical replicates were assessed per treatment group except the study reported in supplementary figure 1 wherein two technical replicates were used for force analysis. A summary of all media and solutions used for hMMT seeding and culture can be found in **Supplementary Table 2**.

### 1.2.5 Electrical field stimulation of hMMTs

Electrical field stimulation (EFS) was performed on day 12 of differentiation as we previously described [24]. Constructs were stimulated at room temperature (RT) immediately after removal from the incubator. Post movements in recorded videos were analyzed to measure the force and kinetics of hMMT contraction using a custom script. This custom, semi-automated script measures post displacement in recorded videos and generates a time course of post locations for the manual analysis of hMMT contraction kinetics. This study implemented an updated version of the script allowing for the simultaneous, semi-automated analysis of hMMT post displacement and kinetics (2.6 and 2.7). The script can be found here [https://github.com/gilbertlabcode/myoTACTIC].

### 1.2.6 Optimization of the EFS parameters for AB1167 hMMTs

A systematic optimization protocol determined the EFS parameters eliciting peak force from AB1167 hMMTs. Briefly, to select the pulse duration and voltage eliciting peak force for 0.5 Hz frequency, hMMTs were stimulated at pulse durations ranging from 1 ms to 80 ms as the voltage was increased by 1 V increments from 1 to 10 V. An EFS frequency of 0.5 Hz was selected since hMMT contractile strength dropped when the frequency was increased to 1 Hz. Constructs were stimulated to contract 6-7 times at each combination of parameters. Average twitch force was calculated as the difference between minimal and peak force values for contractions 3-5 in the stimulus train. Next, to select the frequency eliciting peak tetanic force at 5 V (determined as optimal; see results), the hMMT force-frequency relationship was evaluated. Maximum absolute force during a 3-second stimulation of constructs was quantified at frequencies from 1 to 100 Hz. Absolute force was assessed in 1 Hz increments from 1 to 10 Hz, and in 5 Hz increments thereafter. Three hMMTs from a single MyoTACTIC portion served as replicates. Since hMMT strength decreases with prolonged time at room temperature (unpublished observation), the MyoTACTIC portion containing technical replicates was rewarmed in the incubator for 10-15 minutes following stimulation of each replicate.

### 1.2.7 Evaluation of contractile force and contraction kinetics

EFS at the optimized conditions was performed to assess the contractile function of control and treated hMMTs. Constructs were stimulated for 6-7 contractions at 0.5 Hz, 5 V, and 80 ms to elicit average peak twitch. Following 30 seconds of rest, they were stimulated for 6-7 contractions at 60 Hz, 5 V, and 5 ms to elicit peak tetanic force. Three hMMTs from a single MyoTACTIC portion served as replicates. For 0.5 Hz stimulus trains, we report the average twitch and kinetics of twitch contraction. Time-to-peak tension (TPT), half-relaxation time (1/2 RT), contraction rate, relaxation rate, duration-at-peak, and full-width at half max have been included. Their values were averaged over contractions 3-5 to correspond to the quantification of average twitch. For stimulation at 60 Hz, we report the maximum absolute force and fatigue resistance of tetanic contraction. The absolute force generated during the second tetanic contraction provided a measure of peak force; the percentage of peak force produced during the sixth tetanic contraction was used to quantify fatigue resistance. The first contraction generated by hMMTs was often irregular in comparison to subsequent contractions, and, thus, was excluded.

### 1.2.8 Myotube width and coefficient of variance analysis

At day 12 of differentiation, control and treated hMMTs were fixed in paraformaldehyde (PFA; Cat. #A11313; Alfa Aesar) for morphological analysis. To visualize myotubes within hMMTs, fixed constructs were immunostained for sarcomeric α-actinin (SAA), phalloidin to visualize f-actin, and Hoechst counterstained to observe nuclei. Confocal images were acquired at 40x magnification using the Fluoview-10 imaging software and an IX83 inverted confocal microscope (Olympus). The complete protocol for hMMT staining and imaging has been described [26]. A summary of all solutions and antibodies used to stain hMMTs can be found in **Supplementary Table 3**.

Confocal images of the SAA channel were utilized to quantifyhMMT average myotube width and coefficient of variance. Analysis of flattened image stacks, facilitated by the Fiji Software (ImageJ, NIH), and calculation of these morphometrics was performed in complete accordance with our previously reported methods [26].

### 1.2.9 Nuclear fusion index and SAA coverage analysis

Confocal images of the SAA and Hoechst channels were utilized to quantifyhMMT nuclear fusion index and SAA coverage. All analysis and calculations were performed in complete accordance with our previously reported methods [26].

### 1.2.10 Automated analysis of Z-line architecture

The Z-line architecture of hMMT sarcomeres was assessed using a computational structural assay developed in MATLAB by Morris *et al.* [27]. In short, this program called ZlineDetection extracts a binary skeleton of z-lines from immunostained images of SAA and f-actin which it then uses for the quantification of z-line architecture parameters. For this analysis, hMMT SAA and f-actin confocal images were processed, and the program was run using the same modifications to author-recommended settings as we previously reported [26]. The mean continuous Z-line length, mean number of Z-lines per image, mean sarcomere length, and Orientation Order Parameter (OOP) were evaluated.

### 1.2.11 Statistical Analysis

Statistical analysis was performed using GraphPad Prism 9.0. For data with multiple groups compared across one independent variable, a one-way ANOVA followed by Dunnett’s multiple comparison test was utilized whereby Healthy IgG was set as the comparator. When the effects of two independent variables across multiple groups were compared, a two-way ANOVA followed by Dunnet’s multiple comparison test was utilized to compare differences between the groups to Healthy IgG. All values are expressed as mean ± standard error of the mean (SEM). Significance was defined as p ≤ 0.05. For EFS experiments, Rout’s outlier test, where the maximum desired false discovery rate (Q) was set to Q = 5%, was used to identify outliers. This resulted in the removal of two data points in the Anti-HMGCR group within the “Tetanus (uN)” dataset in **Figure 1B**. Statistical tests were run following the removal of these outliers. A file containing all raw data has been provided.

**Figure 1.**
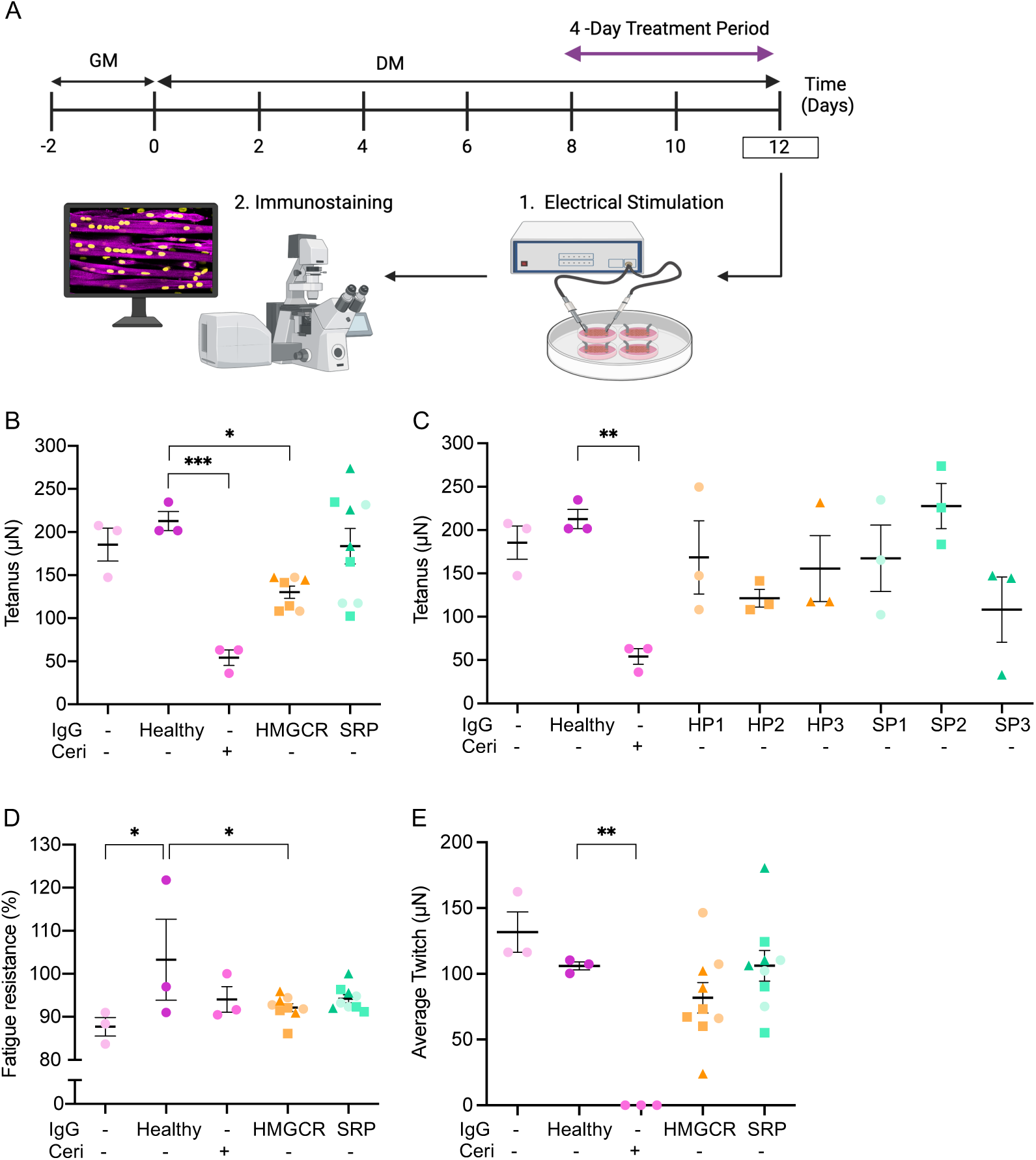
Anti-HMGCR+ autoantibodies reduce hMMT tetanic force. (A) A schematic representation of the hMMT culture timeline and endpoint analyses. hMMTs are cultured in after seeding growth media (GM) and then switched to di<erentiation media (DM) for the next 12 days of culture. On day 8 of di<erentiation, all horse serum containing DM is removed and replaced with heat-inactivated horse serum with or without patient total IgGs. On day 12 of di<erentiation, endpoint analysis is conducted. (B-E) Dot plot of hMMT mean absolute tetanic force with patient groups clustered (B) or separated (C), fatigue resistance during tetanus stimulation (D), and dynamic oscillation of force (average twitch; E). N = 3 patients for anti-HMGCR (anti-HMGC3 Patient 1 [HP1], anti-HMGC3 Patient 2 [HP2], and anti-HMGC3 Patient 3 [HP3]) and for anti-SRP IgGs (anti-SRP Patient 1 [SP1], anti-SRP Patient 2 [SP2], and anti-SRP Patient 3 [SP3]). n = 3 hMMTs per treatment condition.. All values are reported as means ± SEM; * p ≤ 0.05, ** p < 0.01, and *** p < 0.001.

## 1.3 Results

### 1.3.1 Heat-inactivated horse serum does not negatively impact hMMT structure or function

To understand how healthy or patient autoantibodies directly affect muscle health, pre-existing complement from horse serum (HS) was removed by heat inactivation. First, we determined whether heat-inactivated horse serum (HI HS) negatively impacts human skeletal muscle microtissues (hMMTs) by conducting a comparative analysis. Following treatment with HS (standard), HI HS, or HI HS + 25 nM cerivastatin (atrophy-inducing positive control) from day 8 to day 12 of differentiation, hMMTs were subjected to morphometric analysis via immunostaining. Sarcomeric α–actinin and Hoechst 33342 immunostaining at day 12 of differentiation revealed comparable myotube diameter when comparing the HS and HI HS media conditions. As expected, a decline in myotube diameter was observed when cerivastatin was included in the media (**Figure S1A-B**).

To compare the contractile function of hMMTs across different treatment conditions, we first optimized the pulse duration and voltage eliciting average peak twitch at 0.5 Hz and frequency eliciting peak absolute tetanus for hMMTs engineered using the AB1167 immortalized human myoblast cell line [28]. In our optimization, average peak twitch was generated by hMMTs at 80 ms, which we selected as the optimal pulse duration. Electrolysis during EFS above 7 V produced bubbles and caused the media to become murky, precluding optical tracking of post displacement at or above this voltage. Electrolysis did not result from repeated stimulation at 5 V, which we thus defined as our standard voltage. The force-frequency relationship assessed at 5 V demonstrated that maximum absolute force was generated by hMMTs at 60 Hz (**Figure S2**). We consequently selected this as the optimal frequency for evaluation of tetanus force.

hMMTs cultured across the media conditions and then subjected to EFS revealed that force-generating capacity was comparable in hMMTs differentiated with or without HI HS (**Figure S1C-E**). Thus, HI HS was used in all media formulations for the subsequent studies. In the presence of 25 nM cerivastatin, the decline in force-generating capacity at high (**Figure S1C**) or low (**Figure S1D**) frequency EFS did not achieve statistical significance. Additionally, the amplitude of tetanic force did not significantly decline from the first to the last contraction, showing high fatigue resistance (**Figure S1E**). Thus, for all subsequent experiments, we increased the cerivastatin treatment to 75 nM.

### 1.3.2 hMMT EFS reveals reduced tetanic force following exposure to anti-HMGCR aAbs

Total IgGs were extracted from plasmapheresis collected from 3 anti-SRP and 3 anti-HMGCR patients. Clinical features of the patients from which these samples were acquired can be found in **Supplementary Table 4**. To investigate the direct effect of anti-SRP and anti-HMGCR aAbs on hMMT function and structure, we designed an *in vitro* stress test. On day 8 of differentiation, the striated, contractile myotubes in hMMTs were introduced to HI HS DM containing 1 mg/ml total IgG. At the day 12 culture endpoint, hMMTs were subject to EFS to evaluate the function, and then immediately immunostained for morphometric analyses (**Figure 1A**). High-frequency EFS to elicit tetanic contractions revealed a statistically significant decline in absolute contractile force in the presence of anti-HMGCR aAbs, but not anti-SRP aAbs (**Figure 1B**). By visualizing the tetanic contractile force data of hMMTs at the individual HMGCR^+^ and SRP^+^ patient level, we found that all 3 HMGCR^+^ patient samples and 2 SRP^+^ patient samples produced a trended decline in hMMT tetanic contractile force, although none achieved statistical significance (**Figure 1C**). hMMTs from all groups showed strong fatigue resistance, with a slight impairment observed in the (-) Healthy IgG control and the anti-HMGCR groups (**Figure 1D**). During low-frequency EFS, quantification of average twitch revealed a trended decline in contractile force in the presence of anti-HMGCR aAbs that did not achieve statistical significance (**Figure 1E**).

Next, we evaluated contraction kinetics at low-frequency EFS. Quantification of time-to-peak tension (**Figure 2A**), duration-at-peak (**Figure 2B**), half-relaxation time (**Figure 2C**), and full-width at half-maximum (**Figure 2D**) revealed no significant impairments in the presence of anti-HMGCR or anti-SRP aAbs relative to the healthy IgG control. However, the anti-HMGCR aAb treated hMMTs trended towards a decline in the rate of hMMT contraction (**Figure 2E**), and both anti-HMGCR and anti-SRP aAB treated hMMTs trended towards a decline in the rate of relaxation (**Figure 2F**).

**Figure 2.**
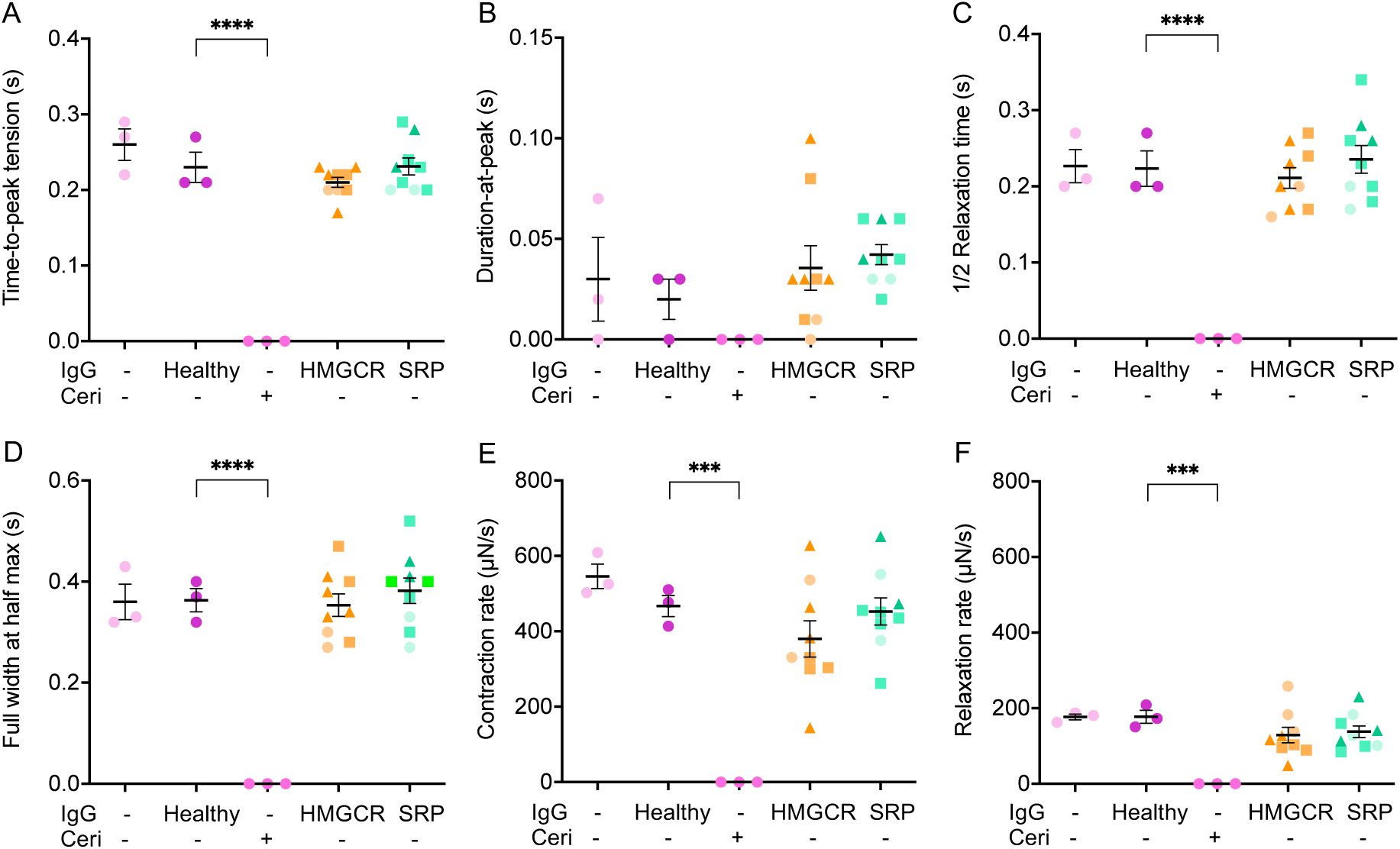
Anti-HMGCR+ autoantibodies impact the rate of hMMT contraction and relaxation. Dot plot quantification of time-to-peak tension (A), duration-at-peak (B), half-relaxation (C), full-width at half-maximum (D), contraction rate (E), and relaxation rate (F). N = 3 for anti-HMGCR and anti-SRP IgGs, n = 3 hMMTs per treatment condition. All values are reported as means ± SEM; * p ≤ 0.05, ** p < 0.01, *** p < 0.001, and **** p < 0.0001.

### 1.3.3 Diminished myotube diameter observed in anti-SRP aAb treated hMMTs

We next examined whether alterations in myotube morphology resulted from treatment with anti-HMGCR and/or anti-SRP autoantibodies following EFS. Sarcomeric α–actinin and Hoechst 33342 immunostaining at day 12 of differentiation revealed the formation of multinucleated, striated, and aligned myotubes in all groups (**Figure 3A**). Upon quantifying the width of myotubes within treatment hMMTs, we found that exposure to total IgGs from anti-SRP^+^ IMNM patients, induced a statistically significant decline in myotube width, as also observed in our cerivastatin positive control for atrophy (**Figure 3B**). An evaluation of myotube diameter frequency uncovered a significant increase in the incidence of smaller myotubes in hMMTs exposed to anti-SRP aAbs compared to healthy IgG control (**Figure 3C**). Mean myotube width in hMMTs treated with anti-HMGCR IgGs trended as smaller than the healthy IgG control, albeit not to statistical significance. Quantification of the coefficient of variance across individual myotubes revealed low variance in myotube width across all groups, with the lowest variance observed in the (-) Healthy IgG control hMMTs (**Figure 3D**). Nuclear fusion index measurements showed that hMMTs across all treatment groups contained a comparable proportion of unfused nuclei suggesting that these autoantibodies do not drive reserve cell expansion or myonuclear accretion (**Figure 3F**). Consistently, quantification of the proportion of sarcomeric α–actinin^+^ structures revealed that treatment groups giving rise to hMMTs with smaller myotubes also contained fewer sarcomeric α–actinin^+^ structures (**Figure 3F**).

**Figure 3.**
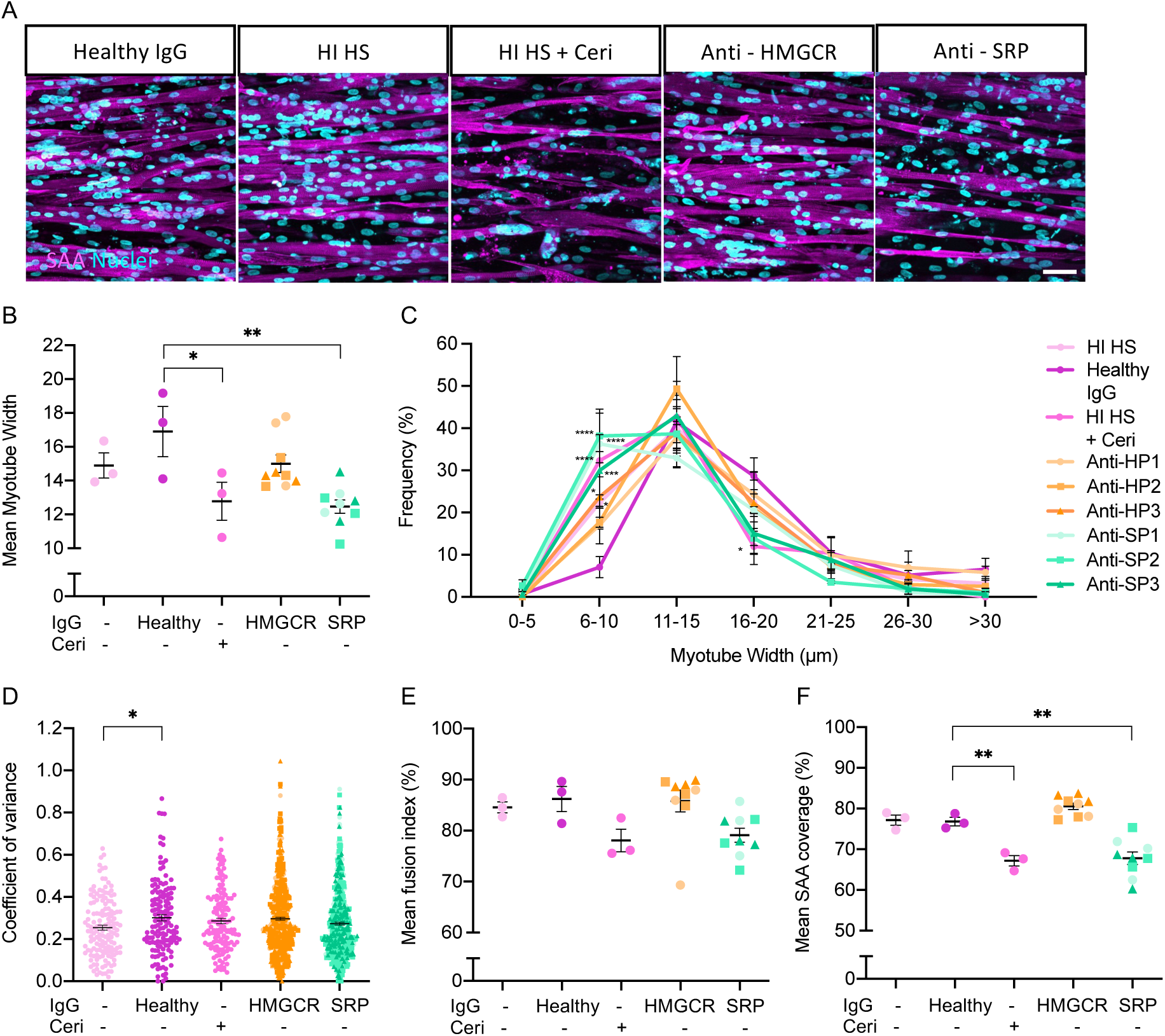
Anti-SRP autoantibodies reduce hMMT myotube diameter. (A) Representative 40x confocal images of myotubes formed in hMMTs within each condition immunostained for sarcomeric α-actinin (SAA, magenta) and counterstained with Hoechst 33342 (cyan). Scale bar = 50 µm. (B) Dot plot showing average myotube diameter for individual hMMTs. (C) Histogram illustrating myotube diameter frequency across treatment conditions. (D) Dot plot of myotube width variation as quantified by the coe<icient of variance for single myotubes. n = 146 for (+) and (-) healthy IgG, n = 138 for Ceri, n = 524 for anti-HMGCR, and n = 550 for anti-SRP. (E) Dot plot of hMMT nuclear fusion index for individual hMMTs. (F) Dot plot of mean SAA positive coverage within flattened confocal stack images for individual hMMTs. (B -F) N = 3 patients for anti-HMGCR and for anti-SRP IgGs. n = 3 hMMTs per treatment condition. All values are reported as means ± SEM; * p < 0.05,** p < 0.01, *** p < 0.001, and **** p < 0.0001.

### 1.3.4 Stable myotube sarcomere organization in aAb treated hMMT following electrical stimulation

Following EFS, we immunostained hMMTs to visualize sarcomeric α–actinin and f-actin (**Figure 4A**, left panel) to analyze the stability of the sarcomere structures following a mechanical load. We have previously shown that in the case of hMMTs produced from a Duchenne muscular dystrophy patient immortalized myoblast line (Δ exons 45-52), striations are sparse or almost eliminated following the same mechanical load that was applied to our IgG-treated tissues (**Figure 4B**)[21]. Here we again implemented an automated analysis tool that enables assessment of sarcomere/z-line architecture. Specifically, we evaluated mean z-lines per image, mean continuous z-line length, mean sarcomere length, and the orientation order parameter (OOP) (**Figure 4**). We found that the mean z-lines per image, mean continuous z-line length, and the mean sarcomere length were similar upon comparing healthy total IgG and anti-SRP+ and anti-HMGCR+ treated hMMTs (**Figure 4C-E**). The OOP, a metric scored out of 1, whereby a higher value signals better sarcomere organization, showed a trended decline in anti-HMGCR treated hMMTs compared to healthy controls, however, this was not statistically significant (**Figure 4F)**.

**Figure 4.**
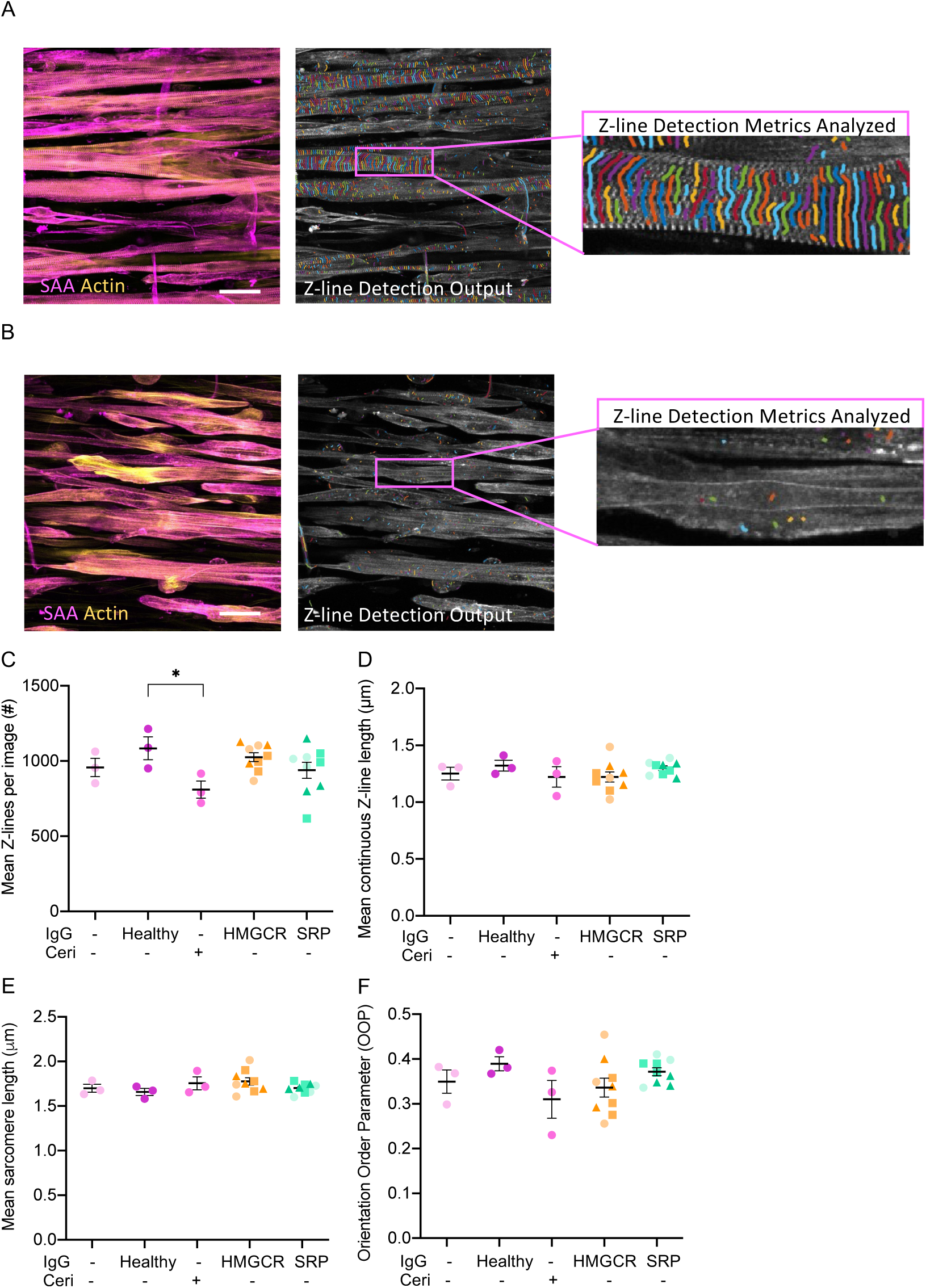
Sarcomere structures are maintained after mechanical load. (A) Left panel shows a representative 40x confocal image of hMMT myotubes post electrical field stimulation immunostained for sarcomeric α-actinin (SAA, magenta) and f-actin (actin, yellow) which corresponds to the input for the ZlineDetection program. Scale bar = 50 µm. Right panel shows a corresponding output from the ZLineDetection program with a further zoom in to illustrate the retention of striations following a mechanical load. (B) Left panel shows a representative 40x confocal image of DMD hMMT myotubes following electrical field stimulation immunostained for SAA (magenta) and actin (yellow). Scale bar = 50 µm. Right panel shows a corresponding output from the ZLine Detection program with a further zoom in to illustrate loss of striations after a mechanical load, as previously quantified [21]. (C-F) Dot plots of mean Z-lines per image (C), orientation order parameter (OOP; D), mean continuous Z-line length (E), and mean sarcomere length (F). N = 3 patients for anti-HMGCR and for anti-SRP IgGs. n = 3 hMMTs per treatment condition. All values are reported as means ± SEM; * p ≤ 0.05.

## 1.4 Discussion

Herein we deliver evidence for direct, but distinct effects of anti-HMGCR+ and anti-SRP+ IMNM patient aAbs on the contractile function of human skeletal muscle. By implementing an advanced skeletal muscle to study IMNM, we were positioned to decouple the secondary immune influences of IMNM-associated aAbs on skeletal muscle function and allowing direct pathogenic influences to be uncovered. We observed that anti-HMGCR+ aAbs induced a decline in hMMT tetanic force, and in the kinetics of tissue contraction and relaxation, without impacting hMMT myotube diameter or sarcomere organization. By contrast, anti-SRP+ aAbs caused myotube atrophy, with limited effects on hMMT contractile force. Our observations suggest distinctions in anti-SRP+ and anti-HMGCR+ IMNM disease pathogenesis that may be important considerations in therapy development.

In our anti-SRP and anti-HMGCR aAb-treated hMMTs, we did not see a positive correlation between myotube width and contractile force (**Figure S3**). Literature suggests that while muscle size and strength are generally positively correlated, the relationship between the two can be complex and depend on various factors [29,30]. Specifically, in IMNM, muscle size and strength can be independently influenced by the degree of inflammation, age of onset, the effectiveness or type of treatment, and individual variation such as comorbidities, to name a few factors. For instance, steroid therapy is considered the first line of therapy for general myositis; however, despite an increase in muscle strength after steroid therapy, researchers observed a significant loss of muscle mass even when there was no underlying disease, suggesting that the therapy itself decreases muscle volume [31].

Clinically, anti-SRP^+^ IMNM patients have more severe disease (atrophy and weakness) than anti-HMGCR^+^ IMNM patients. The direct influences of anti-HMGCR+ patient aAbs on hMMT function we observed, and more subtle impacts of anti-SRP+ patient aAbs, may reflect a distinction in pathogenic mechanism. Alternatively, the inability to observe a decline in contractile force or an alteration of contraction kinetics following treatment with anti-SRP patient aAbs may be due to the spread in hMMT responses we observed, which could reflect patient variability. For example, each patient may produce different polyclonal Abs targeting SRP and HMGCR, differing in target affinity, and resulting in patient-specific distinctions based upon the Ab clones they produce. hMMTs were treated with 1 mg/mL of total IgGs, which is about 10-fold less than IgG concentration in human plasma, which may also explain discrepancies. However, it must be noted that it remains unclear what IgG concentration that muscle fibers are exposed to in vivo. Finally, expanding functional metrics to include assessments of skeletal muscle excitation-contraction coupling [19–21,32,33], assessing myotube necrosis in hMMTs, or introducing aAb treatments earlier culture time-points since SRP and HMGCR were found expressed at the cell membrane during differentiation, may each uncover more dramatic direct pathogenic influences of SRP+ IMNM patient aAbs.

Together our findings corroborate the role of anti-SRP and anti-HMGCR aAbs in IMNM, highlighting humoral mechanisms as targets for IMNM therapies. Additionally, we show that the role of aAbs can be studied *ex vivo*, in an advanced skeletal muscle culture system. It would be informative for future studies to conduct studies specifically using anti-SRP and anti-HMGCR aAbs isolated from the total IgG fraction, to understand whether these disease associated aAbs are indeed responsible for direct pathogenic effects, a finding that would have important implications in the development of CAR-T cell therapies to treat IMNM. Nevertheless, the study herein argues for the use of advanced cell models of skeletal muscle alongside patient-derived tissue and animal studies to distinguish the direct patho-mechanism of aAbs from those involving the complement cascade.

## 1.5 Author Contributions and Acknowledgements

PMG, OB and AM conceived of the project and designed experiments. HL, designed and performed experiments, analyzed data, and prepared figures with assistance from EME and YT. All authors contributed to data interpretation. ZX wrote the semi-automated script for force kinetic analysis which was validated by YT. AM isolated the autoantibodies from patient plasmapheresis and determined their concentration. BT tested autoantibody target recognition. YA, OB, and PMG supervised and funded the research. HL, EME, YT, and PMG wrote the manuscript. All authors edited and approved the manuscript. Schematics were prepared using BioRender.com.

## Supplemental Figure Captions

**Supplementary Figure 1.**
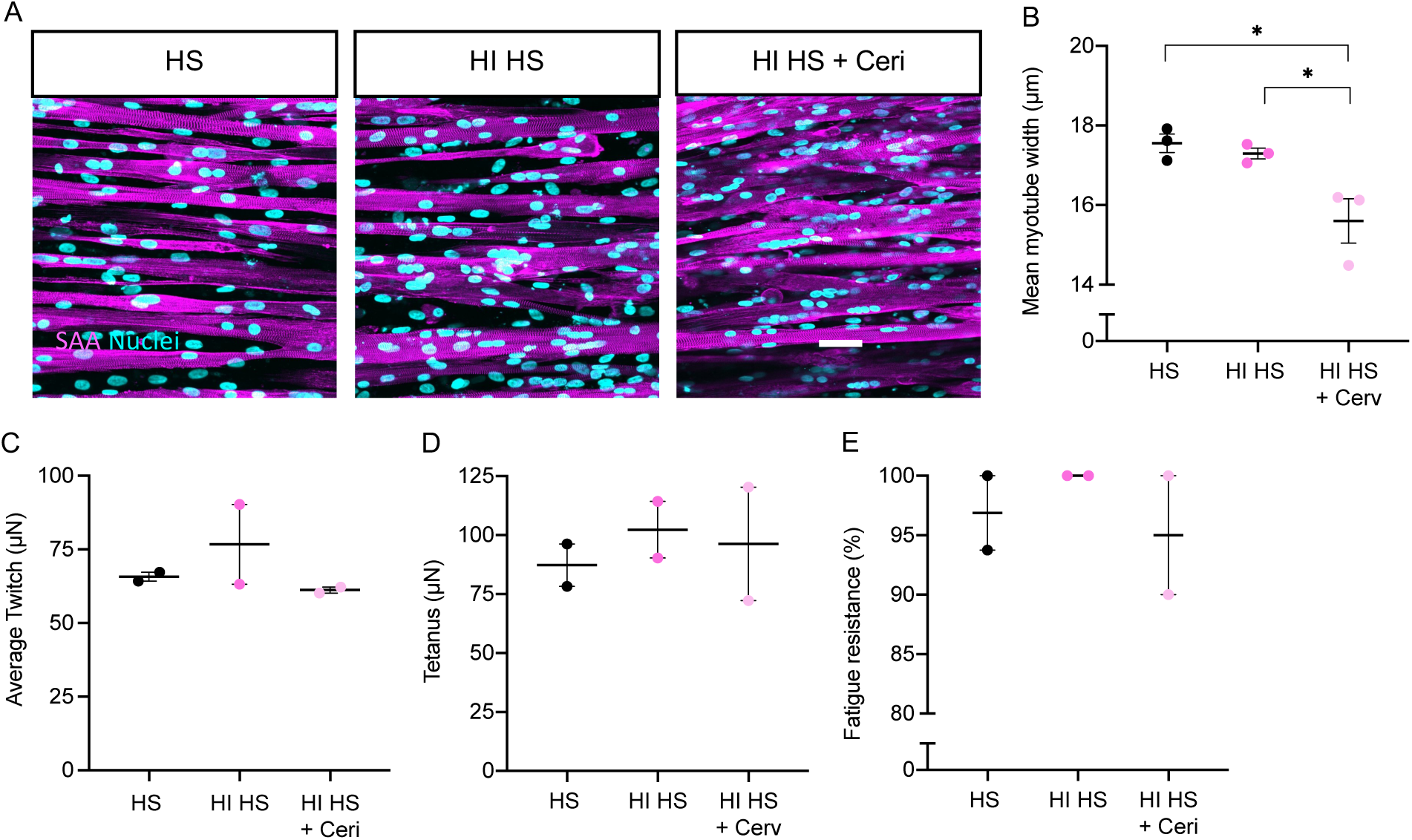
Heat-inactivated horse serum does not affect hMMT myotube size and force. (A) Representative 40x confocal images of myotubes formed in hMMTs within each condition immunostained for sarcomeric α–actinin (SAA, magenta) and counterstained with Hoechst 33342 (cyan). Scale bar = 50 µm. (B) Dot plot showing average myotube diameter quantified for individual hMMTs. n = 3 hMMTs per condition. (C-E) Dot plots showing average twitch (C) and tetanus (D) contractile forces generated by EFS of hMMTs at the optimized stimulation parameters and (E) fatigue resistance over 5 tetanic contractions. n = 2 hMMTs per condition. All values are reported as mean SEM, * p ≤ 0.05. Significance was determined by unpaired one-way ANOVA followed by Dunnet’s multiple comparison test to compare the mean of each column.

**Supplementary Figure 2.**
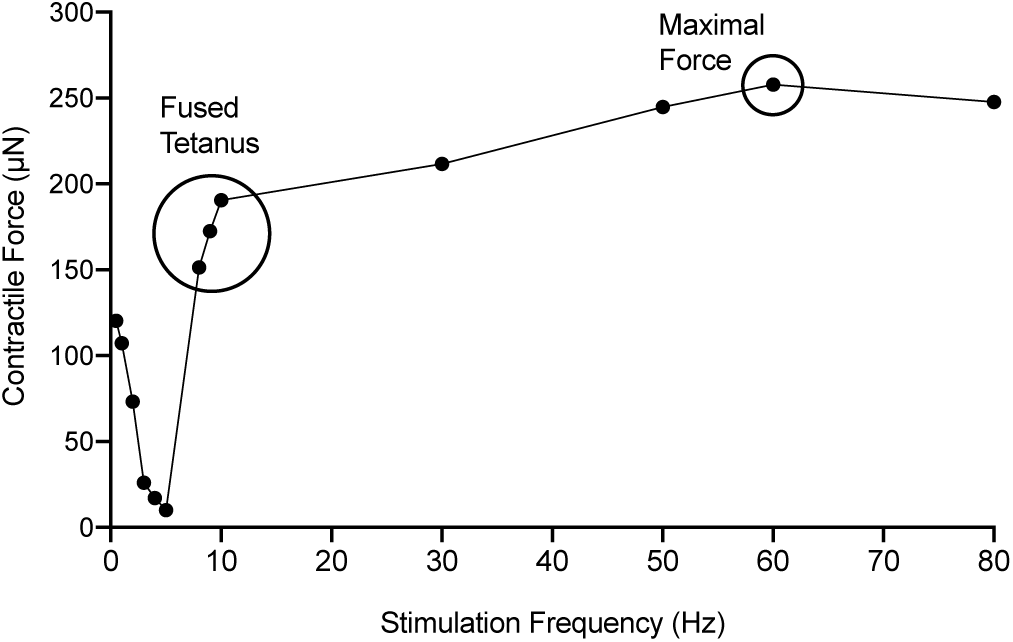
Optimization of electrical field stimulation parameters. Line graph displaying outcomes from optimization of electrical stimulation parameters for high frequency stimulation (tetanus). n = 4 hMMTs.

**Supplementary Figure 3.**
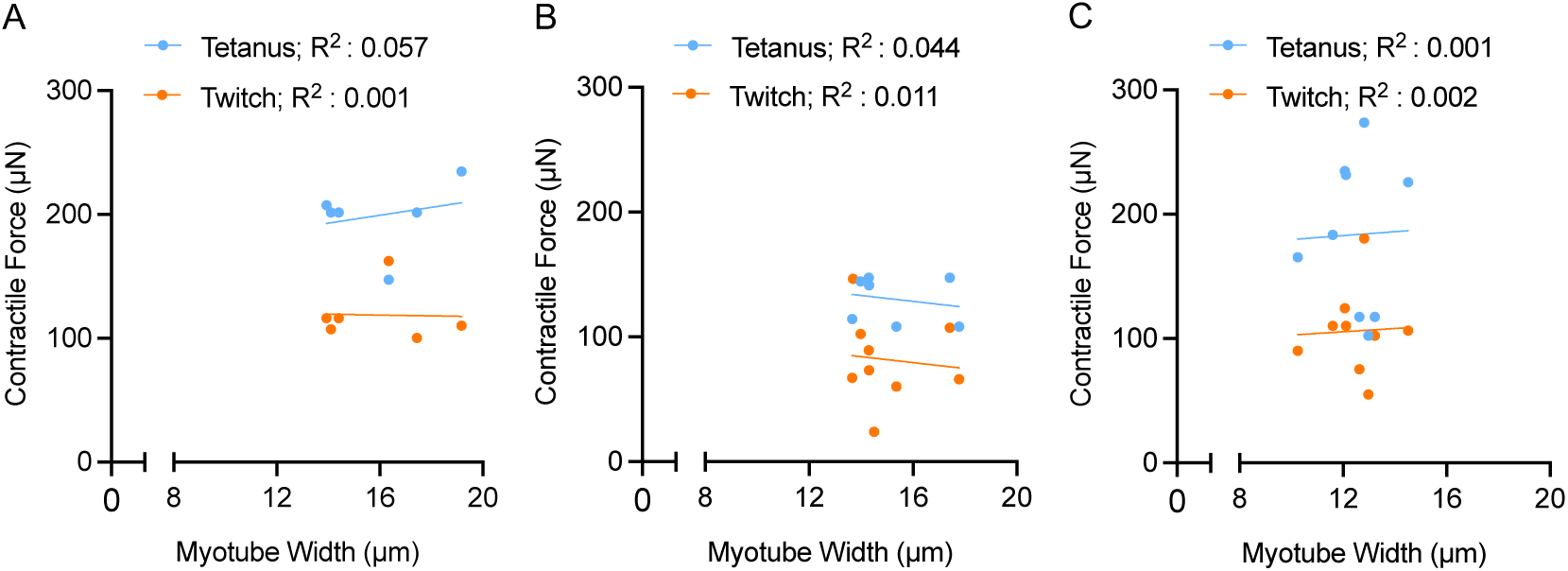
No correlation was found between myotube size and strength. The average myotube diameter (µm) and contractile force (µN) of each (A) Healthy IgG and HI HS, (B) anti-HMGCR and (C) anti-HMGCRP treated hMMT was plotted on the x and y axes, respectively. A linear regression was used to determine the goodness of fit where the R^2^ value revealed no positive correlation between myotube size and hMMT contractile strength.

**Supplementary Table 1.** Concentration of total IgG samples extracted for this study and target recognition.

| Sample ID | Experiment ID | Pathology | Total IgG Conc (mg/ml) | Target Recognition |
| --- | --- | --- | --- | --- |
| SP1 | Anti – SRP<br>Patient 1 | Anti – SRP | 43.613 | 57U |
| SP2 | Anti – SRP<br>Patient 2 | Anti – SRP | 49.09 | >57U |
| SP3 | Anti – SRP<br>Patient 3 | Anti – SRP | 28.02 | 11U |
| HP1 | Anti – HMGCR<br>Patient 1 | Anti – HMGCR | 51.5 | 192.5 CU |
| HP2 | Anti – HMGCR<br>Patient 2 | Anti – HMGCR | 54.4 | 175.7 CU |
| HP3 | Anti – HMGCR<br>Patient 3 | Anti – HMGCR | 48.4 | 60.7 CU |
| Total Plasma Control | Healthy IgG | Negative Control | 47.27 | - |

SRP (signal recognition particle), HMGCR (3-hydroxy-3-methylglutaryl-CoA reductase).
U and CU are recognition units.

**Supplementary Table 2.** Composition of media and solutions used for myoblast maintenance and hMMT production.

| Media/solution | Composition |
| --- | --- |
| hMMT differentiation medium | DMEM<sup>1</sup> (Cat. #11995073, Gibco), 2 % HS (with and without heat inactivation), 2 mg/mL ACA<sup>2</sup> (4 % v/v; Cat. #A2504, Sigma), 10 µg/mL human recombinant insulin (Cat# 91077C, Sigma), 1 % P/S<sup>3</sup> (Cat. #15140122, Gibco) |
| hMMT after seeding growth media | Skeletal Muscle Cell Basal Medium (Cat. #C-23260, Promo-Cell; Heidelberg, Germany), 20 % FBS<sup>4</sup> (Cat. #12483020, Gibco), 1.5 mg/mL ACA (3 % v/v), 1 % P/S |
| Fibrinogen stock solution | 10 mg/mL fibrinogen (Cat. #F8630, Sigma) in 0.9% (wt/v) NaCl solution in water |
| Hydrogel mixture | DMEM (40 % v/v), 4 mg/mL fibrinogen (40 % v/v), Geltrex™ (20 % v/v; Cat. #A1413202, Gibco) |
| Myoblast growth media | Skeletal Muscle Cell Growth Medium kit (Cat. #C-23160, Promo-cell), 15 % FBS, 1% P/S |
| Wash medium | DMEM, 5 % FBS, 1 % P/S |
| Freezing Media | 90% FBS and 10% DMSO<sup>5</sup> |

<sup>1</sup>Dulbecco's Modified Eagle's Medium (DMEM). <sup>2</sup>6-Aminocaproic acid (ACA). <sup>3</sup>Penicillin-Streptomycin (P/S). <sup>4</sup>Fetal bovine serum (FBS). <sup>5</sup>Dimethyl sulfoxide (DMSO).

**Supplementary Table 3.** Solutions and antibodies used for hMMT immunohistochemical staining.

| Solutions | Composition |
| --- | --- |
| Fixing solution | 4 % PFA in D-PBS<sup>1</sup> (Cat. #311-415-CL, Wisent Bioproducts; Saint-Jean-Baptiste, Quebec, Canada) |
| Permeabilization and blocking solution | 10 % GS<sup>2</sup> (Cat. #16210072, Gibco), 0.3 % Triton X-100 (Cat. #TRX777, BioShop; Burlington, Ontario, Canada) in D-PBS |
| Primary antibody solution | Mouse anti-SAA antibody (1:800; Cat. #A7811, Sigma) diluted in blocking solution |
| Secondary antibody solution | Goat anti-mouse IgG conjugated with Alexa-Fluor™ 488 (1:500; Cat. #A-11001, Invitrogen; Waltham, MA, USA ), Phalloidin conjugated with Alexa Fluor™ 568 (1:400; Cat. #A12380, Invitrogen), Hoechst 33342 (1:1000; Cat. #H3570, Invitrogen) diluted in the blocking solution |

<sup>1</sup> Dulbecco's phosphate-buffered saline (D-PBS). <sup>2</sup>Goat serum (GS).

**Supplementary Table 4.** Clinical features and characteristics of the patients from which plasma was acquired.

| Sample ID | Pathology | Sex | Age at Sampling | Treatment at sampling | CK Levels | MMT8 | PGA |
| --- | --- | --- | --- | --- | --- | --- | --- |
| SP1 | SRP | F | 68 | CTC, MTX | 3065 | 90 | 9 |
| SP2 | SRP | F | 27 | Solumedrol, oral corticosteroid, IgIV | 12431 | 97 | 9 |
| SP3 | SRP | F | 26 | CTC, IgIV | 7754 | 105 | 8 |
| HP1 | HMGCR | M | 47 | Zilucoplan, CellCept, IgIV, Cortancyl | 1453 | 94 | 7 |
| HP2 | HMGCR | M | 24 | Cortancyl, Endoxan, IgIV | 269 | 142 | 4 |
| HP3 | HMGCR | F | 34 | CTC | 164 | 102 | 6 |

